# Cytoplasmic localization of ribonucleotide reductase is essential for genome stability

**DOI:** 10.1101/2024.10.29.620917

**Authors:** Christos Andreadis, Israel Salguero, Stephen E. Kearsey

**Author notes:** Spanish National Cancer Research Centre C/ Melchor Fernández Almagro, 3. 28029 Madrid (Israel Salguero).

## Abstract

Ribonucleotide reductase (RNR) is a major determinant of dNTP concentrations and its correct regulation is crucial for genome stability. Active RNR is cytoplasmic but the significance of this localization, away from DNA synthesis, is unclear, particularly as RNR is targeted to the nucleus after DNA damage occurs. We subverted the localization of fission yeast RNR by tagging the large (Cdc22^R1^) and small (Suc22^R2^) subunits with nuclear localization signal sequences. Directing RNR to the nucleus caused a slow S-phase and increased mutagenesis, in spite of predicted constitutive activation of RNR. Defective function of nuclear RNR is also suggested by the formation of Cdc22^R1^ supramolecular filaments and defective RNR reoxidation. In contrast, constitutively nuclear Cdc22 or Suc22 has little effect on DNA damage repair and in fact suppresses *rad3*Δ-sensitivity to DNA damaging agents. We suggest that there is an inadequate flux of substrate through nuclear RNR for S phase, but nuclear targeting of RNR may facilitate DNA damage repair under some circumstances.

## Introduction

Ribonucleotide reductase (RNR) catalyzes the rate-limiting step in *de novo* synthesis of deoxyribonucleotides (dNTPs) from ribonucleotides (NDPs). Correct regulation of RNR is vital for maintaining proper dNTP levels in the cell, ensuring high-fidelity DNA synthesis in replication and repair, thus preserving integrity of both nuclear and mitochondrial genomes (Bebenek et al., 1992, Chabes et al., 2003, Gon et al., 2011, Holmberg et al., 2005, Mathews, 2006, Moss et al., 2010, Pai and Kearsey, 2017). RNR is regulated at a number of different levels, including allosteric control, transcriptional regulation, proteolysis and interaction with small inhibitory proteins (reviewed in (Guarino et al., 2014, Arnaoutov and Dasso, 2014, Holmberg and Nielsen, 2012, Fairman et al., 2011, Nordlund and Reichard, 2006, Cotruvo and Stubbe, 2011)). Upon RNR inhibition, DNA replication slows, leading to activation of the intra-S phase checkpoint, and facilitating preservation of limiting dNTPs (Alvino et al., 2007, Koc et al., 2004). DNA damage leads to upregulation of RNR activity by a variety of mechanisms (Chabes et al., 2003, Holmberg et al., 2005) (reviewed in (Nordlund and Reichard, 2006)), to facilitate DNA synthesis associated with repair. Inability to upregulate dNTP levels by activating RNR during cell proliferation promotes cancer development (Bester et al., 2011), highlighting the importance of RNR in genome stability. Furthermore, inhibitors of RNR such as hydroxyurea, clofarabine and gemcitabine have been successfully employed in chemotherapy of numerous types of cancer (Bonate et al., 2006, Shao et al., 2006). It has been found that the most common base lesion in replicating yeast and mammalian cells is misincorporation of riboNMPs, instead of dNMPs. Thus, another consequence of reduced dNTP levels is likely to be increased riboNMP incorporation into replicated DNA (Yao et al., 2013), which can lead to genome instability and, if riboNMP removal is defective, autoimmunity (Reijns et al., 2012, Clark et al., 2011). While low levels of RNR activity and corresponding low levels of dNTPs are detrimental to genome stability, high dNTP levels also affect DNA replication by reducing proofreading fidelity (Mathews, 2006, Buckland et al., 2014) and by stimulating the activity of low fidelity translesion synthesis (TLS) polymerases (Prakash and Prakash, 2002, Sabouri et al., 2008). During the cell cycle, dNTP levels are down-regulated outside of S phase and in nonproliferating cells, and in mammalian cells this down-regulation may be a strategy to inhibit viral replication (Allouch et al., 2013). Furthermore, dNTP synthesis has been found to be facilitated through promotion of MBF gene transcription via Set2-dependent H3K36 methylation (Pai et al., 2017) and that maintaining dNTP homeostasis is critical in preventing replication failure in response to CDK-induced replication stress (Pai et al., 2019).

Eukaryotes use a class I RNR, which is composed of large (R1) and small (R2) subunits. R1 has the catalytic site and two sites for allosteric regulation whereas R2 provides a tyrosyl free radical that is transferred to the catalytic site to effect substrate reduction. The stoichiometry of active RNR is unclear; a tetramer composed of two large and two small subunits appears to be the minimally active complex (reviewed in (Eklund et al., 2001)), but evidence suggests that higher order complexes may constitute the active form (Fairman et al., 2011, Hofer et al., 2012). A conserved aspect of active RNR in eukaryotes is its cytoplasmic localization, with transport or diffusion of deoxyribonucleotides to sites of DNA synthesis in mitochondria or the nucleus respectively (Pontarin et al., 2008, Engstrom et al., 1984), but the biological significance of this localization, away from sites of DNA synthesis, is unclear. In mammalian cells, R1, R2 and p53R2 subunits are constitutively cytoplasmic in the absence of DNA damage (Pontarin et al., 2008). However, in *S. cerevisiae* and *S.pombe,* R1 is cytoplasmic or pancellular (Liu et al., 2003), while R2 is regulated by cytoplasmic localization, in that it is nuclear outside of S phase, but exported to the cytoplasm during S phase or after DNA damage to activate RNR. In *S. cerevisiae*, nuclear import of R2 is promoted by Dif1, and R2 is anchored in the nucleus by association with Wtm1. Activation of the DNA damage checkpoint leads to Dif1 phosphorylation and degradation, and release of R2 from Wtm1 allows formation of active R1-R2 complexes in the cytoplasm (Wu and Huang, 2008, Lee et al., 2008). In fission yeast, upon DNA damage, Spd1 is downregulated, following ubiquitylation by CRL4-Cdt2, abolishing its interaction with R1 (Cdc22^R1^) in the cytoplasm and allowing RNR activation (Holmberg et al., 2005, Hakansson et al., 2006, Liu et al., 2005, Liu et al., 2003, Salguero et al., 2012).

In addition to cell cycle changes in RNR subunit localization, there have been several reports suggesting that, after DNA damage, RNR is localized to sites of DNA repair (Niida et al., 2010a, D’Angiolella et al., 2012, Hu et al., 2012, Zhang et al., 2009, Tanaka et al., 2000). This may facilitate repair synthesis, particularly at cell cycle stages when dNTP levels are depressed, in a manner which does not require a global increase in dNTP levels. Both large (hRRM1) and small subunits (hRRM2 and p53R2) are translocated to the nucleus after DNA damage (Tanaka et al., 2000, Xue et al., 2003). This accumulation requires Tip60-RRM1 interaction and abolishing this interaction sensitizes cells to DNA damage (Niida et al., 2010a). Furthermore, Cyclin F and RRM2 physically interact and colocalize to the nucleus in G2, and cyclin F-mediated RRM2 degradation seems to prevent genome instability (D’Angiolella et al., 2012). In addition to RNR, thymidylate kinase is targeted to sites of DNA repair (Hu et al., 2012). In tobacco, the R1a subunit is transiently relocalized from the cytoplasm to the nucleus upon DNA damage possibly reflecting a role in RNR transcriptional activation (Lincker et al., 2004). Thus, nuclear RNR might have additional functions such as regulation of transcription, especially since RNR-Tip60 complex can be formed even in non-DNA damage conditions (Niida et al., 2010a). However, these observations are controversial as other reports have failed to detect nuclear RNR in mammalian cells after DNA damage, suggesting cytoplasmic dNTPs can diffuse sufficiently fast into the nucleus in a manner that is not rate limiting for DNA synthesis or repair (Pontarin et al., 2008). Importantly, in *S. cerevisiae* it has been demonstrated that MCM helicase physically interacts with RNR and some of its regulators such as Dun1 kinase occur within small subpopulations of RNR-MCM, independently of their subcellular locations (Yáñez-Vilches et al., 2024). When these interactions are impaired, Rad52 release from DNA repair foci is disrupted, leading to hypermutagenesis. This highlights the non-canonical role which RNR can play in preserving the genetic stability of the cell (Yáñez-Vilches et al., 2024).

In this investigation we study the importance of regulating RNR by Suc22^R2^ cellular localization and the significance of the cytoplasmic localization of active RNR by directing Cdc22^R1^ and Suc22^R2^ subunits to the nucleus. Preventing colocalization of Cdc22^R1^ and Suc22^R2^ during S phase and after DNA damage impedes DNA replication but has relatively little effect on DNA damage sensitivity. Forcing the nuclear colocalization of both Cdc22^R1^ and Suc22^R2^ subunits has little effect on dNTP levels and DNA damage sensitivity but leads to a suboptimal S phase and increased mutation rate and has a different phenotype from when both Cdc22^R1^ and Suc22^R2^ subunits are cytoplasmic. Our results suggest that the highly conserved cytoplasmic localization of RNR in eukaryotic cells has evolved to optimise the dNTP supply for S phase, but nuclear RNR may not be detrimental and, in some circumstances may facilitate, DNA repair.

## Results

### Redirecting fission yeast RNR activity to the nucleus

We subverted the normal cytoplasmic localization of active RNR by directing single or both large Cdc22^R1^ and small Suc22^R2^ subunits to the nucleus. The endogenous *suc22* and *cdc22* genes, expressed from their native promoters, were modified so that the protein products have N-terminal NLS-GFP sequences (Fig.1A, right), and the cellular distribution of these proteins was compared to fluorescently-tagged versions of the proteins lacking NLSs (Fig. 1 A, left). As previously reported (Liu et al., 2003), Suc22^R2^ is normally nuclear for most of the cell cycle and shows redistribution to the cytoplasm after inhibiting DNA synthesis with HU (Fig.1B, top panels). After tagging with NLS-GFP of Suc22 ^R2^ (henceforth referred to as NLS-Suc22^R2^), the subunit is constitutively nuclear both in exponentially growing cells and after treatment with HU (Fig. 1B, bottom panels). Cdc22^R1^ lacking an NLS is constitutively pancellular (Fig. 1C, top panels, and (Liu et al., 2003)), but after growth in HU, a high proportion of cells show aggregation of Cdc22^R1^ into cytoplasmic foci (Fig. 1C, top panels). After NLS-GFP tagging of Cdc22 ^R1^ (henceforth referred to as NLS-Cdc22^R1^), the protein is nuclear, but rather than a uniform pan-nuclear distribution, shows accumulation in filaments which can be up to about 3 μm long, spanning the nucleus (Fig. 1C, and 1E), and the distribution does not change significantly after arrest in S-phase following treatment with HU (Fig. 1C, bottom panels). In *S. cerevisiae*, RNR has been reported to form large supramolecular foci (Noree et al., 2010), and we speculate that the fission yeast nuclear environment or higher local concentration of Cdc22 ^R1^ may in some way enhance the interaction between Cdc22^R1^ subunits, leading to the formation of filament structures.

**Figure 1.**
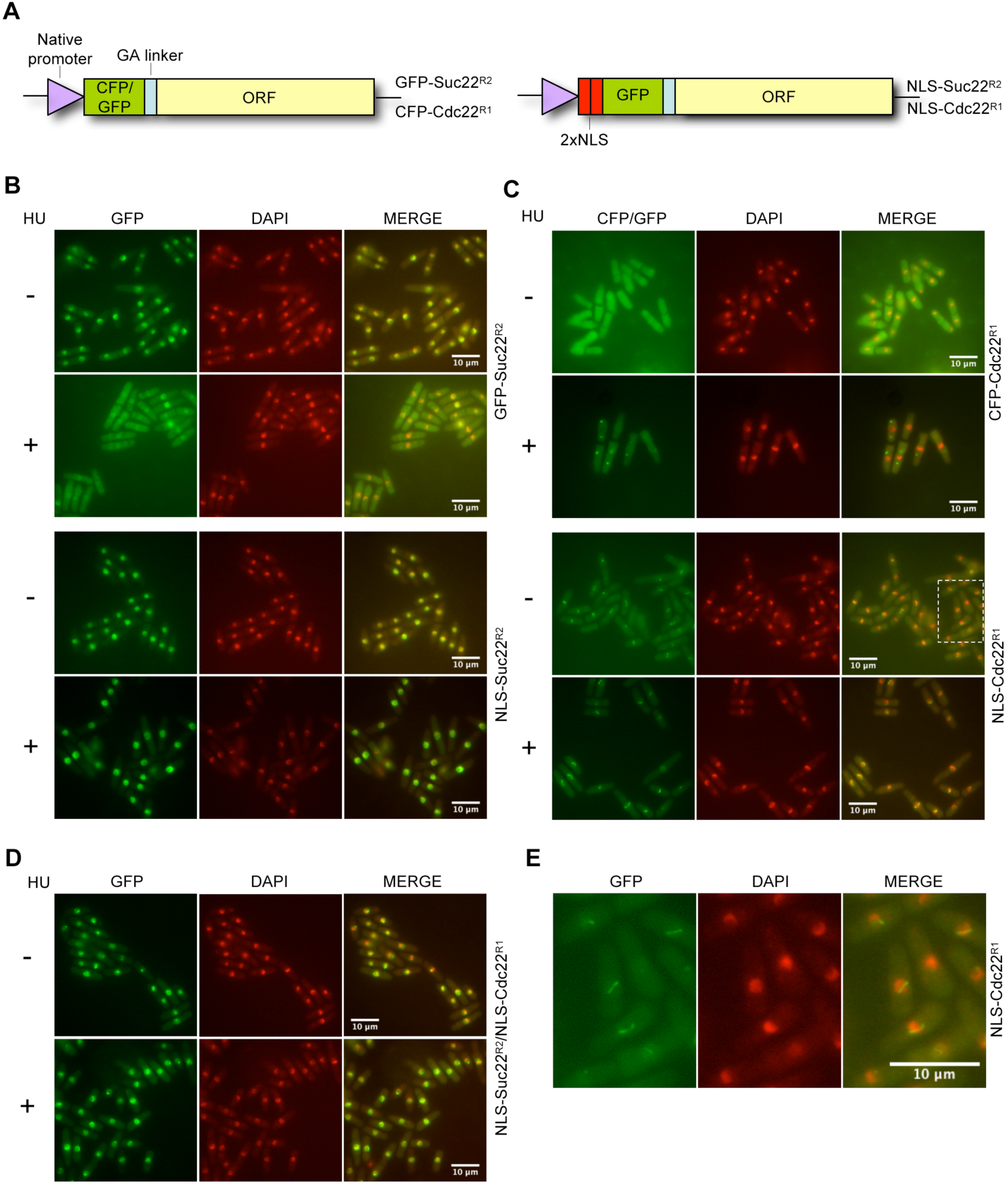
Redirecting RNR subunits to the nucleus. (A) Schematic representation of RNR gene construction expressing GFP or NLS-GFP N-terminal tagged proteins. Genes were regulated by their native promoters. (B-E) Localization of RNR in exponentially growing cells was examined by fluorescence microscopy. HU treatment (12mM) was for 2 hours. Cells were fixed using methanol/acetone, but similar results were obtained by imaging live cells (not shown). (B) GFP-Suc22^R2^ (2342) and NLS-Suc22^R2^ (3167) localization in the absence or presence of HU (top and bottom panels, respectively). (C) CFP-Cdc22 (2344) and NLS-Cdc22^R1^ (3276) localization in the absence or presence of HU (top and bottom panels, respectively). (D) NLS-Cdc22^R1^ NLS-Suc22^R2^ (3277) localization in the absence or presence of HU. (E) Magnification of area inside dashed rectangle from panel C (NLS-Cdc22^R1^ cells).

We also constructed a strain where both CFP-Cdc22^R1^ and GFP-Suc22^R2^ subunits are localized to the nucleus (Fig. 1D). These cells show similar nuclear distributions of proteins in exponential growth and after treatment with HU, with nuclear filaments, suggesting that the supramolecular structures seen with NLS-Cdc22^R1^ alone are not affected by NLS-Suc22^R2^.

Each cycle of reduction of NDP by RNR generates a disulphide bond in the R1 active site. It has been proposed that this is reduced by a C-terminal pair of cysteines of another R1 subunit, before the newly formed disulphide bond at the C-terminus is reduced by thioredoxin or glutaredoxin (Zhang et al., 2007), so we investigated whether Cdc22^R1^ filaments seen in NLS-Cdc22^R1^ might be due to aberrant intersubunit disulphide bond formation. Analysis on reducing gels showed that protein levels are affected by nuclear localization, with Cdc22^R1^ levels decreased and Suc22^R2^ levels increased compared to non-NLS versions (Fig. 2A) Co-localization of both subunits to the nucleus did not affect protein levels compared to strains where single subunits were constitutively nuclear. Western blot analysis under non-reducing conditions showed additional bands with wild-type Cdc22^R1^ at a position corresponding to a Cdc22 ^R1^ dimer, which is resolved by treatment with a reducing agent (Fig. 2B). With the NLS-Cdc22^R1^ NLS-Suc22^R2^ strain, a high molecular weight smear is seen which is partially resolved by reduction, although in the strain where only NLS-Cdc22^R1^ is present, this smear is barely visible (Fig. 2B). Thus, although this suggests that some unresolved disulphide bond formation involving nuclear Cdc22^R1^ and/or Suc22^R2^ subunits leads to formation of high molecular weight complexes, this is unlikely to be responsible for the supramolecular structures seen by fluorescence microscopy, which are equally prominent in the NLS-Cdc22^R1^ NLS-Suc22^R2^ and NLS-Cdc22^R1^ strains.

**Figure 2.**
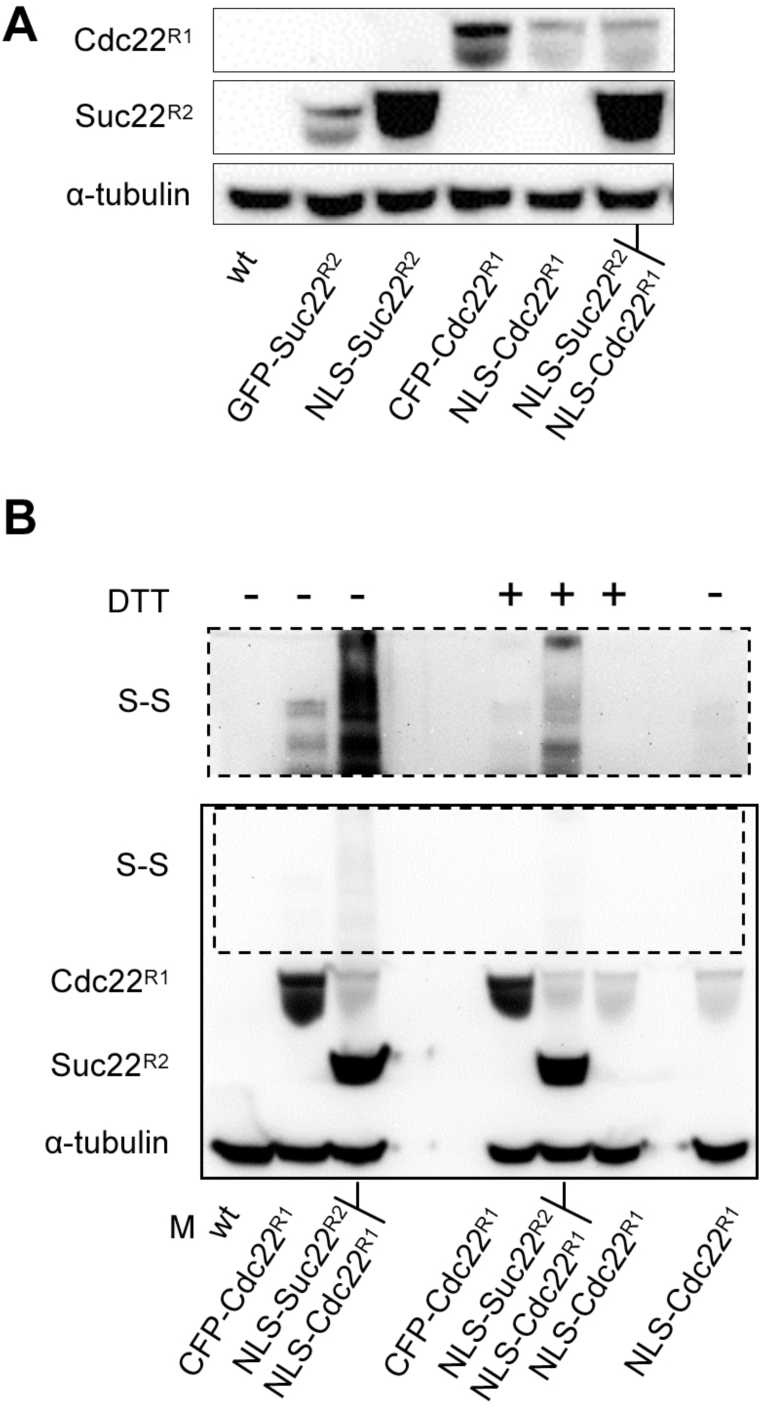
Western blot analysis of nucleus-targeted RNR. (A) Western blot analysis of RNR in NLS-and non-NLS RNR strains (strains used from left to right: 138, 2342, 3167, 2344, 3276, 3277). (B) Western blot analysis of non-reduced RNR samples in NLS- and non-NLS RNR strains in the presence or absence of DTT reducing agent in the sample buffer, as indicated. Possible Cdc22 dimers, due to S-S bond formation, are shown in the top panel, which represents a longer exposure of the bottom dashed rectangle. Strains used from left to right: 138, 2344, 3277, 2344, 3277, 3276, 3276. In both panels tubulin is used as a loading control.

### Effect of targeting RNR subunits to the nucleus on dNTP levels

We used HPLC to determine how dNTP concentrations are affected by nuclear localization of RNR subunits (Fig. 3A). N-terminal tagging of Cdc22^R1^ with CFP had no effect on dNTP levels compared to wild-type (wt), while similarly tagged Suc22^R2^ showed approximately 50% of wt dNTP level suggesting that the tag, while tolerated, may interfere with RNR function.

**Figure 3.**
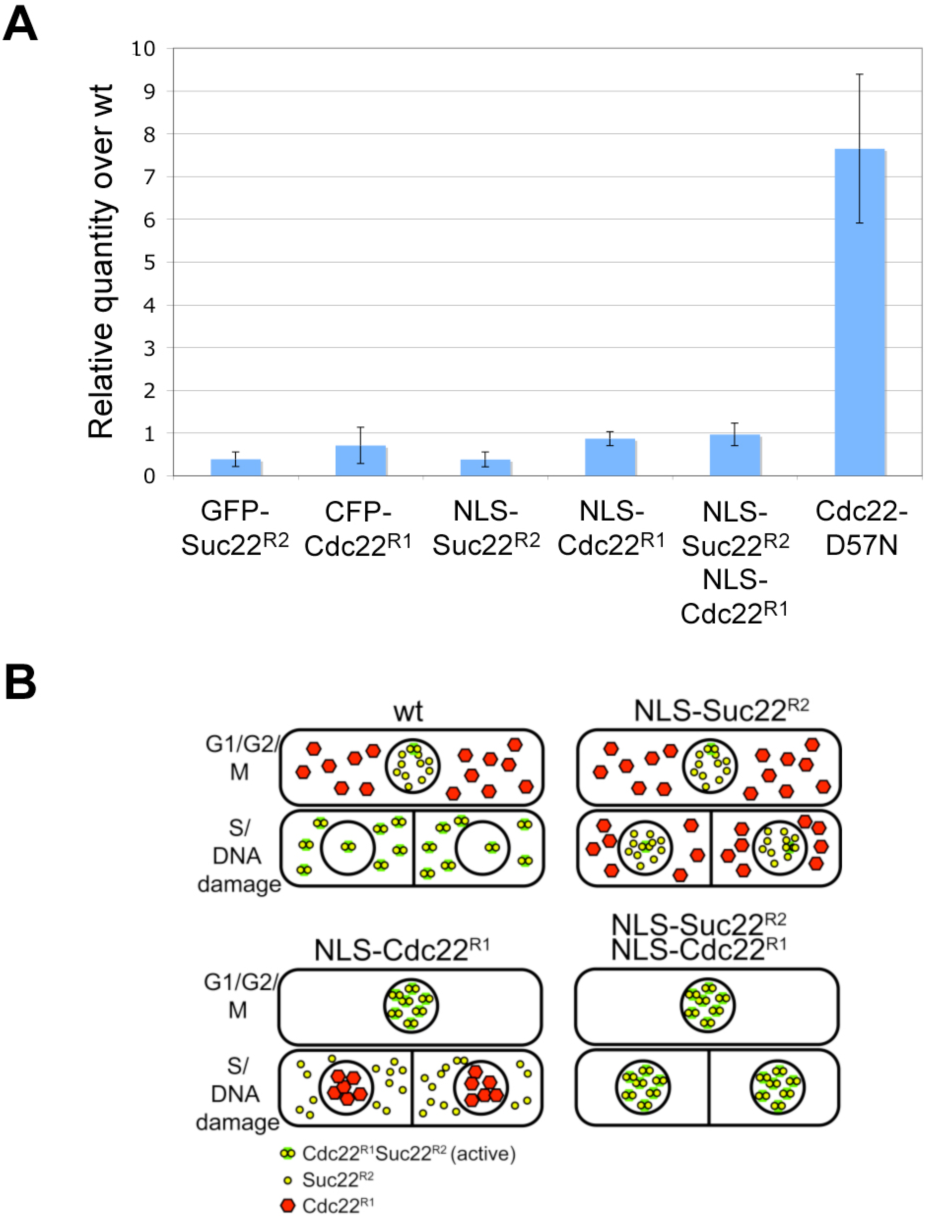
Effect of RNR nuclear localization on dNTP levels. (A) dNTP levels were measured by HPLC; values shown are averages of three determinations; error bars show the standard deviation. Strains used were 2342, 2344, 3167, 3276, 3277 and 3285 (left to right). (B) Predicted localization of Cdc22^R1^ and Suc22^R2^ in wt and NLS-targeted strains in G1/G2/M phases and in S phase/after DNA damage.

Constitutively nuclear localization of Cdc22^R1^ would be predicted to increase RNR activity for most of the cell cycle, when Suc22^R2^ is also localized in the nucleus (Fig 3B). Conversely, during S phase, reduced RNR activity would be predicted, since Suc22^R2^ is relocalized to the cytoplasm but Cdc22^R1^ will remain nuclear. Reduced RNR activity would also be predicted from the lower level of Cdc22^R1^ in the NLS strain. These opposing effects may account for the measured dNTP level in the NLS-Cdc22^R1^ strain, which is approximately wt (Fig. 3A).

Constitutive localization of Suc22^R2^ to the nucleus would be predicted to lead to low levels of RNR activity, since outside of S phase Suc22^R2^ is nuclear anyway, but preventing Suc22^R2^ nuclear export during S phase would block formation of active cytoplasmic RNR. However, the measured dNTP concentration in the NLS-Suc22^R2^ strain is about the same as in the control GFP-Suc22^R2^ strain. It is possible that the higher level of Suc22^R2^ protein seen in the NLS strain may compensate for the reduction in RNR activity expected from prevention of nuclear export.

Constitutive localization of both Cdc22^R1^ and Suc22^R2^ to the nucleus would be predicted to lead to increased RNR activity, since both Suc22^R2^ and Cdc22^R1^ are in the same compartment for the entire cell cycle, rather than being in separate compartments outside of S phase (Fig 3B). Thus, an elevation of dNTP levels might be expected but the levels are approximately wt. This could be partly due to the reduction in Cdc22^R1^ levels seen when the subunit is nuclear-targeted. For comparison, dNTP levels are much more significantly elevated in a Cdc22-D57N mutant, where the dATP allosteric downregulation of RNR activity is abolished (Fig 3A and (Fleck et al., 2013)). Taken together, these results suggest that blocking the regulation of Suc22^R2^ by nuclear export and forcing the nuclear localization of active RNR have minor effects on dNTP levels, perhaps due to other compensatory regulatory mechanisms, and the confounding effects of changed RNR subunit levels.

### Constitutively nuclear RNR inhibits growth and S phase

We examined growth rate and S phase progression to determine if they are affected by constitutively nuclear RNR subunits. CFP-Cdc22^R1^ and GFP-Suc22^R2^ tagged strains show similar growth rates to wt (Fig 4A). However, strains where one or both RNR subunits are NLS-tagged show a slight reduction in growth rate, consistent with a slightly bigger cell size in NLS-cdc22 NLS-Suc22, NLS-Cdc22 and CFP-Cdc22 (Fig 4B, left panel). Analysis of DNA contents of exponentially growing cells showed that the NLS-Cdc22^R1^ and NLS-Cdc22^R1^ NLS-Suc22^R2^ strains exhibit slow S phase progression in that there is a <2C shoulder on the histogram, although the NLS-Suc22^R2^ strain shows no such defect (Fig 4B, right panel).

**Figure 4.**
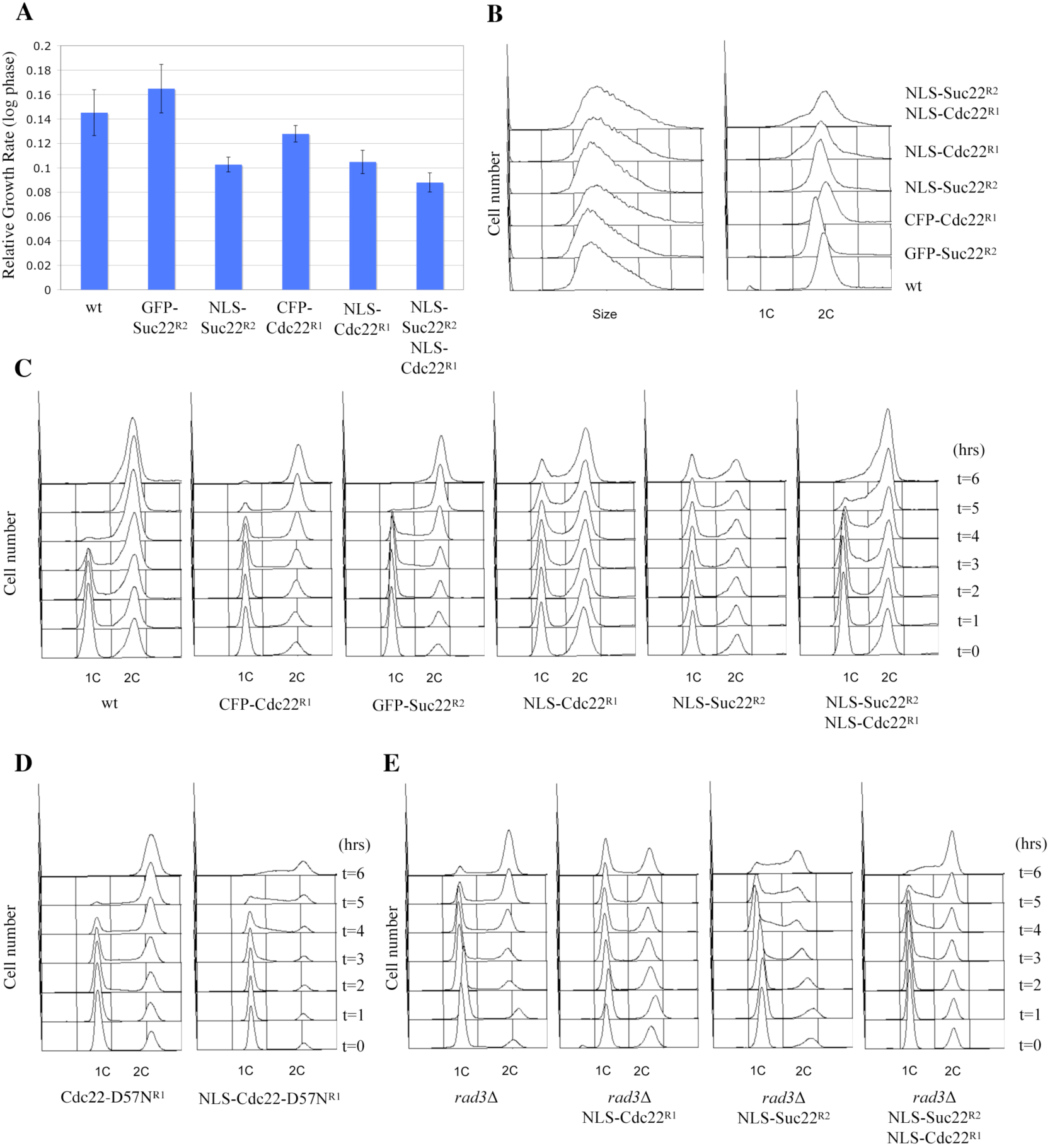
Cell cycle analyses of NLS-RNR strains. (A) Relative growth rates of each strain (strains used were 138, 2342, 3167, 2344, 3276, 3277; left to right). (B) Size (forward scatter) and DNA content of log phase strains after flow cytometric analysis (strains used were 138, 2342, 2344, 3167, 3276, 3277; bottom to top). (C-E) Analysis of S phase execution in cells released from minus-nitrogen block. Cells were released from a G1 block and DNA contents were analysed by flow cytometry at the times shown. Strains used were from left to right in (C) 2299, 3164, 3549, 3328, 3327, 3329; in (D) 3576, 3581; in (E) 3602, 3584, 3620, 3613.

Since it can be difficult to see S phase defects by analysing unsychronised cultures, we performed flow cytometry analysis of S phase progression in cells released from a nitrogen-starvation induced G1 block (Fig. 4C). S phase in the wt was completed 5 hours after release. Both CFP-Cdc22^R1^ and GFP-Suc22^R2^ strains showed a slightly slower S phase, but S phase was complete by 6 hours. In contrast, cells expressing NLS-Cdc22^R1^ and NLS-Suc22^R2^ came out of the G1 block more slowly, and a G1/S phase population was still present 3-6 hours after release, especially for the NLS-Suc22^R2^ strain. This slow S phase can be accounted for by the predicted lowered dNTP levels in S phase, as RNR subunits are directed to different compartments. Although the dNTP measurements show no dramatic reduction, these were made on unsynchronised cells, so significant changes in dNTP levels in S phase may be obscured since this is a relatively short fraction of the cell cycle. However, the fact that these strains are viable indicates that sufficient RNR activation is occurring in spite of NLS tagging. So, in the NLS-Suc22^R2^ strain, there is perhaps sufficient nuclear Cdc22^R1^, and in the NLS-Cdc22^R1^ strain, a pool of Suc22^R2^ may remain nuclear or traffic to the nucleus during S phase, in both cases providing an adequate dNTP supply for DNA synthesis.

Targeting both RNR subunits to the nucleus resulted in a faster S phase compared to when either Suc22^R2^ or Cdc22^R1^ were nucleus-targeted, but still a substantial number of cells were in S phase at 4-6 h following G1 block and release. Paradoxically, RNR activity would be expected to be high in the NLS-Cdc22^R1^ NLS-Suc22^R2^ strain, so the slow S phase is unexpected.

To analyse whether the slow S phase seen in nucleus-targeted RNR strains is due to reduced dNTP levels, we looked at the effects of deleting the Spd1 RNR inhibitor, which results in an ca. two-fold increase in dNTP levels (Fleck et al., 2013). We found that depletion of Spd1 does eradicate the <2C profile seen in the NLS-Cdc22^R1^ strain consistent with the notion that low dNTP levels in this strain are constraining S phase (Supplementary Fig. S1). We saw no effect of deleting Spd2, consistent with a study showing that this protein is not an RNR inhibitor (Vejrup-Hansen et al., 2014). However, *spd1*Δ did not suppress the slow S phase seen in the NLS-Cdc22^R1^ NLS-Suc22^R2^ strain (Supplementary Fig. S1), arguing that the slow S phase shown in this strain is not due to a dNTP insufficiency. As an alternative way of increasing dNTP levels, we investigated whether the slow S phase of an NLS-Cdc22^R1^ strain could be suppressed by the Cdc22-D57N mutation, which prevents allosteric downregulation of RNR by dATP and in *S. pombe* results in a 6-12 increase in dNTP levels (Fleck et al., 2013). A slight delay to S phase is seen in a Cdc22-D57N mutant compared to wt (Fig. 4D) but an NLS-Cdc22-D57N^R1^ strain completes S phase faster than an NLS-Cdc22^R1^ strain, consistent with dNTP insufficiency being a cause of the slow S phase (Fig 4D).

### Slow S phase seen in strains with constitutively nuclear RNR is not dependent on DNA damage checkpoint activation

Abnormally low dNTP levels can cause activation of the DNA replication checkpoint, which can slow S phase (Kim and Huberman, 2001; Santocanale and Diffley, 1998). We therefore investigated whether inactivation of this checkpoint pathway can alleviate slow DNA replication seen when RNR subunits are constitutively nuclear. *rad3*Δ strains, expressing NLS-Suc22^R2^, NLS-Cdc22^R1^ or NLS-Cdc22^R1^ NLS-Suc22^R2^ were arrested in G1 by nitrogen starvation and released into S phase (Fig. 4E). We found that in the *rad3*Δ control strain, S phase was similar to wt and completed in 6 h following G1 block and release. In contrast, depletion of Rad3 in NLS-Cdc22^R1^, NLS-Suc22^R2^, and NLS-Cdc22^R1^ NLS-Suc22^R2^ strains had little effect on S phase kinetics compared to the *rad3*^+^ strain. Thus, these observations indicate that the S phase delay is not due to activation of the DNA replication checkpoint.

### Constitutively nuclear RNR causes enhanced resistance to DNA damaging agents when the DNA damage checkpoint is inactivated

Elevation of dNTP levels in response to DNA damage promotes DNA repair in yeasts, and failure to increase nucleotide levels leads to defects in repair of DSBs and other lesions. Therefore, defective DNA damage repair in the nuclear RNR strains would suggest low dNTP supply problems, while increased resistance to DNA damage could indicate high dNTP levels, possibly enhanced by the nuclear localization of active RNR. Strains expressing either NLS-Cdc22^R1^ or NLS-Suc22^R2^ show similar sensitivity to a number of DNA damaging agents (camptothecin, MMS, UV, 4-NQO) to control strains, possibly with a slightly increased sensitivity to DNA Double Strand (DSB) formation by bleocin/bleomycin (Fig. 5A). GFP tagging of Suc22^R2^ causes increased sensitivity to HU, but this is suppressed by nuclear localization, possibly because Suc22 levels are higher in NLS-Suc22 (Fig 2A). Thus, preventing colocalization of RNR subunits in the same compartment does not appear to have a dramatic negative effect on DNA repair efficiency, implying that the local dNTP supply for repair pathways is adequate. Cells expressing both NLS-Cdc22^R1^ and NLS-Suc22^R2^, with predicted constitutively active nuclear RNR, show similar sensitivity to MMS, UV and HU to wt cells with a marginal increase in resistance to 4NQO (Fig. 5A), compared to GFP-Suc22^R2^ and CFP-Cdc22^R1^. In contrast, the NLS-Suc22^R2^ NLS-Cdc22^R1^ strain shows slight sensitivity to bleocin/bleomycin (Fig. 5A), suggesting that DSB repair is compromised by nuclear RNR. Thus, despite the effects on S phase, NLS-Suc22^R2^ NLS-Cdc22^R1^ strain does not show marked changes in DNA damage sensitivity.

**Figure 5.**
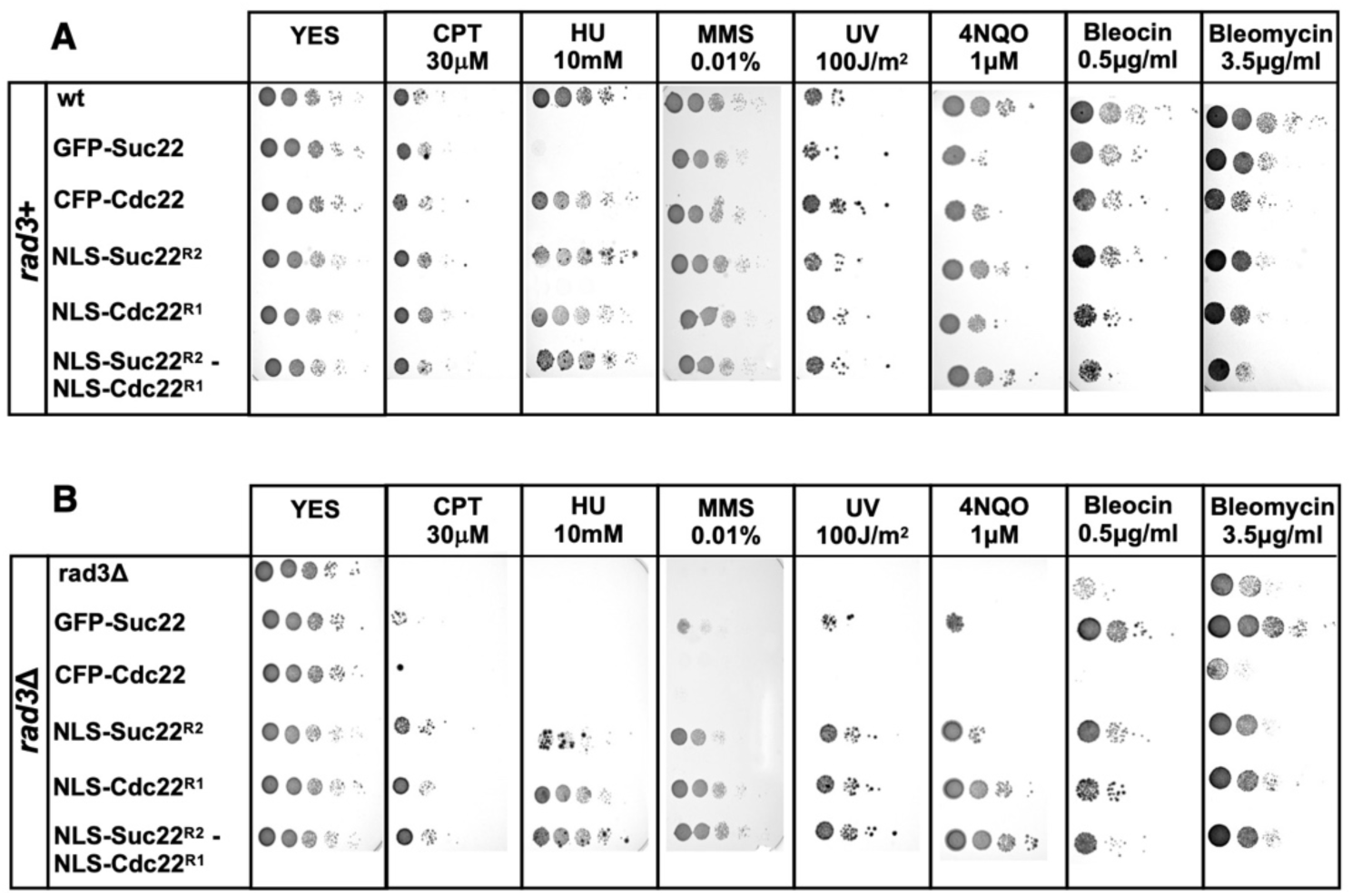
Analysis DNA damage sensitivity and checkpoint activation in NLS-RNR strains. (A-B) Analysis of DNA damage sensitivity of NLS-RNR strains in *rad3*+ (A) or *rad3*Δ (B) background. Serial dilutions of strains were spotted on YES plates containing additives as shown, or were irradiated with UV-C. Representative images of triplicate analysis is shown. Strains used from top to bottom were in (A) 138, 2342, 2344, 3167, 3276, 3277, and in (B) 3406, 3423, 3407, 3428, 3408 and 3409.

A more striking difference is seen when DNA damage sensitivity is compared in a *rad3*Δ background. *rad3*Δ cells are DNA damage sensitive in part because Spd1 is not degraded and RNR is not up-regulated. Indeed, *rad3*Δ DNA damage sensitivity can be suppressed by deleting Spd1 to increase RNR activity (Moss et al., 2010). The rad*3*Δ NLS-Cdc22^R1^ and NLS-Suc22^R2^ strains show clear resistance to DNA damage induced by 4NQO, UV, MMS camptothecin and bleocin/bleomycin, as well as resistance to HU (Fig. 5B). The *rad3*Δ NLS-Suc22^R2^ strain shows higher resistance to DNA damage, although not as high as seen with NLS-Cdc22^R1^ strains (Fig. 5B). Possibly, an increased level of Suc22^R2^ seen in the NLS-Suc22 ^R2^ strain or the nuclear localization of active RNR accounts for this effect. Since Spd1 is predominantly cytoplasmic (Borgne and Nurse, 2000), sequestering NLS-Cdc22^R1^ in the nucleus may reduce inhibition by Spd1.

We measured dNTP levels in *rad3*Δ strains to correlate these with effects on DNA damage sensitivity. In the *rad3*Δ strain, DNA damage by MMS or camptothecin results in a less than two-fold increase in the dNTP level (Supplementary Fig. S2), consistent with previous studies, and smaller than the increase seen in wt cells (Moss et al., 2010). In the NLS-Cdc22^R1^ and NLS-Cdc22^R1^ NLS-Suc22^R2^ strains, an approximately three-fold increase in dNTP levels is seen after DNA damage, consistent with the observation that nuclear targeting of RNR suppresses the *rad3*Δ-associated dNTP defect. In the NLS-Suc22^R2^ strain, the dNTP level is more markedly stimulated after DNA damage compared to the *rad3*Δ control, but less than that seen in the NLS-Cdc22^R1^ and NLS-Cdc22^R1^ NLS-Suc22^R2^ strains, consistent with the predicted effect of constitutive nuclear localization of Suc22^R2^ on dNTP levels. Taken together these results indicated that the DNA repair defects of a *rad3*Δ strain are suppressed by nuclear localization of RNR, but in a wt background there is not marked repair enhancement or deficiency, in contrast to the replication defects seen in these strains. We speculate that the dNTP demands of DNA repair, but not S phase, can be met by nuclear RNR as repair makes less of a demand on dNTP supply compared to replication, and nuclear RNR is compromised for maximum activation.

### Constitutively nuclear RNR results in an increased mutation rate

Correctly balanced levels of dNTPs are critical for high fidelity of DNA replication in S phase. For instance, high levels of dNTPs are associated with reduced fidelity of proofreading and low levels may increase the mutation rate by promoting riboNMP incorporation or polymerase switching (Yao et al., 2013, Holmberg et al., 2005). Thus, we assessed mutation rates in strains with nuclear RNR to see if this aspect of genome stability was disturbed. Using assays based on acquisition of canavanine resistance and reversion of an *ade6* mutation, we detected the most dramatic (two-to five-fold) increase in mutation rates for the NLS-Cdc22^R1^ NLS-Suc22^2^ strain (Fig. 6A, P=0.01; Fig. 6B, P∼0.05). This strain is predicted to have active RNR in the nucleus in S phase and measured dNTP levels are similar to wt. It is difficult to explain this result on the basis of globally elevated dNTP pools. For comparison, the Cdc22-D57N mutant causes a comparable increase in the *ade6-485* reversion mutation rate, but has an eight-fold increase in dNTP level (Fig. 3A), and *spd1*Δ doubles the dNTP level but has no effect on the reversion rate (Fleck et al., 2013). One interpretation of these results is that a local nuclear increase in dNTP levels is contributing to the reduced fidelity of DNA replication and this is not detectable in unsynchronized cells. The NLS-Cdc22^R1^ and NLS-Suc22^R2^ strains show a smaller increase in mutation rate in the CanR assay, which is consistent with the prediction that these strains are compromised in upregulating dNTP levels in S phase.

**Figure 6.**
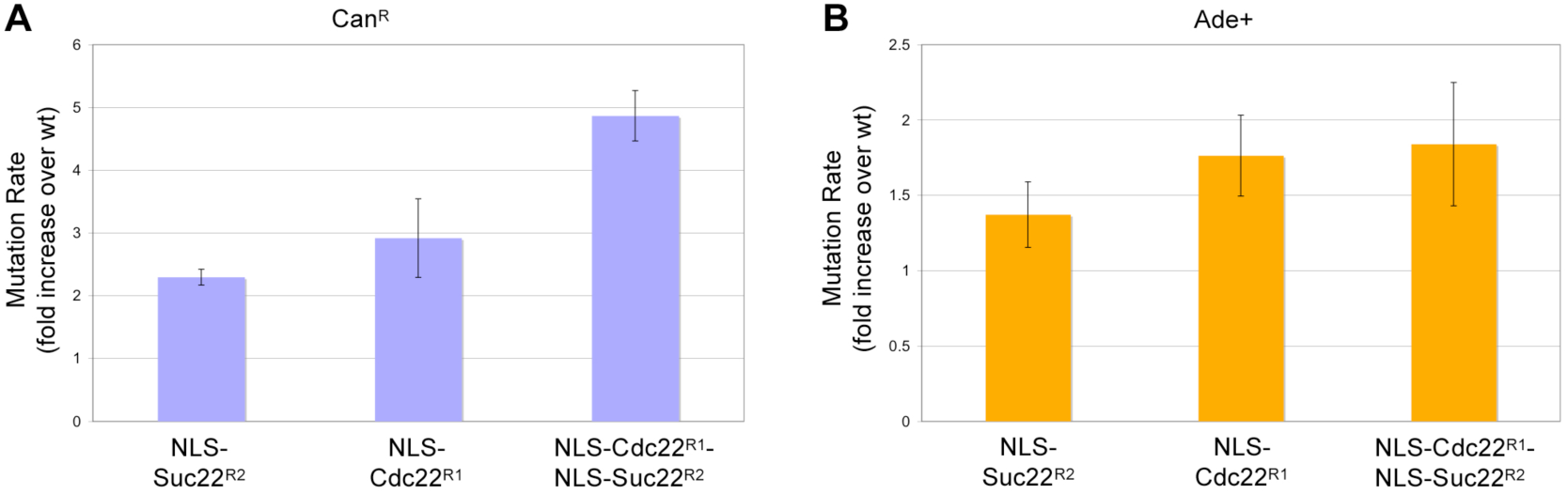
Nuclear RNR causes increased mutation rates. Mutation rates (fold increase over wt) of NLS-RNR mutants were determined by performing fluctuation assays using (A) mutation to canavanine resistance (Can^R^) and (B) reversion of an *ade6-M210* mutation. Error bars represent the standard deviation after 3 determinations. Strains used were 138, 3167, 3276, 3277 and the wt control was 138. For Can^R^, the approximately five-fold mutation rate increase in NLS-Cdc22^R1^ NLS-Suc22^R2^ over wt is significant (P=0.01, 2-tailed equal variance t-test). For the reversion of the *ade6-M210* mutation, the mutation rate change of the NLS-Cdc22^R1^ NLS-Suc22^R2^ over the wt is increased by approximately two-fold (P∼0.05, 2-tailed equal variance t-test).

## Discussion

In this study we have addressed the biological significance of the cytoplasmic localization of active RNR and cellular relocalization of the R2 subunit as a regulatory mechanism. Our main findings are that, despite the normal relocalization of Suc22^R2^ to the cytoplasm during S phase for RNR activation, constitutive targeting of both subunits to the nucleus does not increase dNTP levels, as assessed from asynchronous cells. However, RNR protein levels, S phase progression, and genome stability are misregulated. The presence of CFP-Cdc22^R1^ filaments is a striking consequence of directing this subunit to the nucleus. When Cdc22^R1^ or Suc22^R2^ subunits are directed in the nucleus, S phase execution is slowed. When both RNR subunits are targeted in the nucleus, S phase execution is faster compared to the single NLS-Cdc22^R1^ or NLS-Suc22^R2^ strains, but still slower compared to a wild-type S phase. None of these effects is dependent on activation of the DNA damage checkpoint. Targeting single or both RNR subunits to the nucleus suppresses DNA damage sensitivity in a *rad3*Δ checkpoint deficient mutant. Finally, targeting Cdc22^R1^, Suc22^R2^, or both RNR subunits to the nucleus increases the spontaneous mutation rate. These results indicate that regulation of RNR by subcellular relocalization of Suc22^R2^ is an important but not essential aspect of dNTP control. Furthermore, nuclear localization of RNR is suboptimal for normal S phase execution but apparently not DNA repair. This result may explain why active RNR is evolutionarily conserved in the cytoplasm.

### Altered levels of RNR subunits and filament formation of nuclear-targeted Cdc22^R1^

Unexpectedly, targeting of Cdc22^R1^ and Suc22^R2^ to the nucleus resulted in a reduction and elevation, respectively, of protein levels, and in addition a high proportion of cells show filaments of Cdc22^R1^. Both Cdc22^R1^ and Suc22^R2^ are stable proteins (Christiano et al., 2014), so the increase in Suc22^R2^ levels is likely to be due to increased expression. Suc22^R2^ is upregulated by DNA damage and heat shock (Fernandez Sarabia et al., 1993, Harris et al., 1996) via a mechanism that involves cytoplasmic polyadenylation of its mRNA (Saitoh et al., 2002). This upregulation may also be triggered by constitutive nuclear localization of Suc22^R2^. With regard to the nuclear filaments of Cdc22^R1^, this is unlikely to be an artifact due to GFP tagging, as filaments are not seen with GFP-tagged Cdc22^R1^ in the cytoplasm. The artifactual aggregation was not an issue in a large study of GFP-tagged proteins, which also reported cytoplasmic R1 foci in *S. cerevisiae* (Christiano et al., 2014). Thus, it is possible that filament formation may lead to the reduced protein level of NLS-Cdc22^R1^, and S phase defects. Filament formation does not require Suc22^R2^, and we do not observe abundant non-reduced Cdc22^R1^ in a strain only expressing NLS-Cdc22^R1^, suggesting that disulphide bond formation is not involved. In *E. coli*, supramolecular assemblies of gemcitabine-inhibited RNR have been reported, where R1_4_R2_4_ rings interlock to form large catenated structures (Zimanyi et al., 2012). By performing *in vitro* assays of small-angle X-ray scattering (SAXS) and electron microscopy, Ando et al. (2016) observed that at high ATP concentrations, the ATP-induced hexamer of the nucleotide-binding subunit of human RNR can interconvert with higher-order filaments. Furthermore, in *Bacillus subtilis*, filament formation of class Ib RNR has also been reported by using similar methods (Thomas et al., 2019). In our case, it is unclear why these structures are only seen with NLS-Cdc22^R1^, but it is possible that a higher concentration of the subunit in the nucleus compared to the cytoplasm could be responsible.

### Effect on S phase and DNA repair by inhibiting RNR subunits relocalization to the same compartment

In wt cells, relocalization of Suc22^R2^ to the cytoplasm in S phase is part of the mechanism to activate RNR and dNTP production for S phase, and a similar relocalization occurs after DNA damage. In *S. pombe*, S phase levels of dNTPs are about two-fold higher than during the rest of the cell cycle (Hakansson et al., 2006), but cell cycle relocalization of Suc22^R2^ only partly contributes to this mechanism. Targeting only one of the two RNR subunits to the nucleus is predicted to subvert the activation of RNR in S phase by keeping the subunits in different cellular compartments. In the case of NLS-Cdc22^R1^, Suc22^R2^ will be relocalized to the cytoplasm during S phase, while with the NLS-Suc22^R2^ strain, Suc22^R2^ nuclear export in S phase will be blocked (Fig. 3B). Thus, in both these strains, the dNTP supply may be compromised in S phase, and this is consistent with the observed slow S phase and an increased mutation rate. Although measurement of the dNTP level in the NLS-Cdc22^R1^ strain shows no difference from wt, the increase in RNR activity outside of S phase (when both NLS-Cdc22^R1^ and Suc22^R2^ are nuclear) may mask a reduction in S phase.

After DNA damage, Suc22^R2^ is also relocalized to the cytoplasm where it can form an active complex with Cdc22^R1^. This reactivation of RNR is predicted to be blocked in the NLS-Suc22^R2^ or NLS-Cdc22^R1^ strain, with a detrimental influence on DNA repair. In reality, effects on DNA repair are minor, possibly with an increased sensitivity to DSB-inducing agents, consistent with a requirement for upregulation of dNTPs to facilitate DSB repair (Moss et al., 2010).

### Effect on S phase and DNA repair by constitutive localization of RNR to the same compartment

In contrast to strains where a single RNR subunit is targeted to the nucleus, targeting both subunits would not be expected to compromise dNTP supply during S phase and after DNA damage, since this mirrors the wt situation with active RNR in the nucleus instead of the cytoplasm. Consistent with this prediction, dNTP measurements show a similar level to wt. Unexpectedly, this strain still shows a slow S phase and an increased mutation rate compared to wt. The effect on S phase is not suppressed by *spd1*Δ, suggesting that it is not due to reduced dNTP levels, but rather a consequence of having active RNR in the wrong compartment. In terms of sensitivity to DNA damaging agents, the NLS-Suc22^R2^ NLS-Cdc22^R1^ strain shows an increased sensitivity to DSB-inducing agents, and a slightly increased resistance to 4-NQO, suggesting that the efficiencies of DSB repair and NER are differentially affected by nuclear RNR, but the effects are relatively minor. Similar to *spd1*Δ, NLS-Suc22^R2^ NLS-Cdc22^R1^ also suppresses the DNA damage sensitivity of *rad3*Δ, presumably as there is no requirement for checkpoint activation to allow upregulation of dNTP supply. In all, these results suggest that the normal cytoplasmic localization of RNR is an important factor for normal S phase.

If the effect on S phase is not due to reduced dNTP levels, what could be the explanation? High levels of dNTPs have been reported to delay S phase in mammalian cells (Franzolin et al., 2013), and also in *S. cerevisiae* where the loading of Cdc45 during replication initiation is affected (Chabes and Stillman, 2007). Although dNTP levels are apparently normal in NLS-Cdc22^R1^ NLS-Suc22^R2^ cells, it is possible that a privileged dNTP pool generated by nuclear RNR does not equilibrate quickly with the bulk dNTP pool across the cell, and a locally high dNTP level has inhibitory effects on S phase. An early study suggested that nuclear and cytoplasmic dNTP levels are not equal (Skoog and Bjursell, 1974) and the observed localization of RNR to sites of DNA damage in mammalian cells is assumed to facilitate DNA repair by providing a local dNTP supply, as if the diffusion rate of dNTPs is a limiting factor if RNR is active only in the cytoplasm. It would be interesting to see how constitutively low dNTP levels such as the ones observed after Wee1 inactivation leading to increased dNTP demand and replication stress through CDK-induced firing of replication origins (Pai et al., 2019) would affect RNR subunits localization.

A second possibility is that maintaining dNTP concentration at an optimal level during S phase is less efficient due to an inadequate flux of substrate through nuclear RNR. The pool of dNTPs in S phase is only sufficient for a small fraction of genome synthesis in S phase and RNR must be efficiently upregulated to provide adequate dNTPs (Chabes et al., 2003; Hakansson et al., 2006; reviewed in Niida et al., 2010b). Inefficiencies in this buffering process might not be manifested in measurements of dNTP levels in asynchronous cells. This is less likely to be a factor in DNA repair, as there is not such a demand on nucleotide supply. This explanation could be related to the reactivation of RNR, since every cycle of reduction leads to the formation of a disulphide bond in the R1 subunit, which must be reduced for the next cycle of reduction (Zahedi Avval and Holmgren, 2009), and conceivably this process is less efficient in the nucleus than the cytoplasm. In support of this notion, we observe high molecular weight complexes on nonreducing gels in cells expressing NLS-Suc22^R2^ NLS-Cdc22^R1^, which are partially resolved by reduction. Alternatively, the NDP substrate for RNR, or the conversion of dNDPs to dNTPs may be more limiting in the nucleus than in the cytoplasm.

Overall, these results stress that the rate and fidelity of DNA replication are better supported by cytoplasmic rather than nuclear RNR, which is implied by the evolutionary conservation of RNR localization. RNR activity in the cytoplasm followed diffusion into the nucleus may constitute a buffering mechanism that avoids local high concentrations of dNTPs that might occur if RNR were active in the nucleus. In contrast, nuclear RNR is adequate and in some circumstances beneficial for DNA repair. Subverting the cellular regulation of RNR by blocking the colocalization of RNR subunits in the same compartment is not lethal but is suboptimal for DNA synthesis. This suggests that while the regulation of RNR by cytoplasmic relocalization is an important regulating factor, it is moderated by other regulatory mechanisms, a conclusion also reached by analysis of Spd1 mutants showing that the correct compartmentalization of Suc22^R2^ is not required for RNR regulation (Nestoras et al., 2010). Simultaneous regulation of RNR by a wide range of regulatory mechanisms may together guard against stochastic deviations in dNTP concentrations, analogous to overlapping mechanisms that prevent reinitiation at origins, and contribute to genome stability during DNA replication.

## Materials and Methods

### Strains, constructs and growth conditions

Supplementary Table S1. summarizes the strains used in this study and Supplementary Table S2 lists the primers used for the construction and verification of the strains. Yeast media and genetic techniques were as described (Alfa et al., 1993).

NLS-GFP tagging of the proteins was performed after cloning the gene sequence in frame with 2xNLS, GFP, and (GA)_5_ sequences in a pFA6a plasmid and transforming it into *S. pombe* cells. More specifically, for the pFA-suc22 promoter-NLS-GFP-suc22(N-ter) plasmid, GFP was amplified from the plasmid pSMHG2+ with the primers 1056F and 1057R. Subsequently, both the PCR product and the plasmid pFA6a-suc22 promoter-VC155-suc22 were digested with PacI and BamHI. After gel purification, the insert and the plasmid were ligated and integrated in strain 138 after linearization with XhoI. Yeast transformants were checked by PCR using primers 1112F/996R. For the pFA6a-cdc22promoter-NLS-GFP-cdc22(N-ter) plasmid, the NLS-GFP fragment was obtained by PCR as above. The cdc22 promoter fragment was amplified using primers 952F and 1063R, using genomic DNA as template. The PCR products were annealed and extended to generate the cdc22promoter-NLS-GFP-GA5 fragment. This fragment and the plasmid pFA6a-KanMX6-cdc22 promoter-VN173-cdc22 were digested with BglII and BamHI, gel-purified and ligated. The pFA6a-cdc22promoter-NLS-GFP-cdc22(N-ter) plasmid was then digested with XcmI and integrated in the strain 138. Yeast transformants were checked by PCR using primers 1112F/997R.

*rad3*Δ versions of the strains were constructed by PCR-based gene targeting using KS-ura4 plasmid, as previously described (Bahler et al., 1998), using primers 1306F and 1307R; yeast transformants were screened using primers 1308F and 1309R. *spd1*Δ versions of the strains were constructed by PCR based gene targeting, using the KS-ura4 plasmid in order to get the *spd1Δ::ura4^+^* versions of strains, using primers 1304F and 1305R; yeast transformants were screened using primers 1305R and 536F. Similarly, *spd2*Δ versions of the strains were constructed by PCR-based gene targeting using pClon-Hyg1 plasmid in order to get the *spd2*Δ::hyg1+ versions of strains, using primers 1310F and 1311R; yeast transformants were screened using primers 1312F and 1313R. *spd1*Δ *spd2*Δ versions of strains were obtained by using the PCR-based gene targeting method, using pClon-Hyg1 plasmid (primers 1310F/1311R for *spd2* gene deletion) and transforming *spd1*Δ::ura4+ strains in order to get the *spd1*Δ::ura4+ *spd2*Δ::hyg1+ strains; yeast transformants were screened using primers 1312F/1313R and 1310F/1313R. NLS-Cdc22-D57N versions of the strains were constructed by homologous recombination, after transforming Cdc22-D57N strain (strain number 3285) with a linearized (with XcmI) plasmid pFA6a-cdc22promoter-NLS-GFP-cdc22(N-ter) (see above). The yeast transformants were checked by PCR amplification of the *cdc22* fragment (primers 1112F and 989R) and by digesting the PCR amplicon with DdeI (restriction site generated by the mutation).

### Microscopy

Cells were grown in YE3S to exponential phase, concentrated by centrifugation and fixed with methanol/acetone before being mixed with YE3S mountant (1.2% LMT agarose in YE3S, 0.1mM N-propyl gallate, to reduce phototoxicity) containing 50ng/ml DAPI DNA dye. When live cells were studied, they were grown in YE3S to exponential phase, before being mixed with YE3S mountant (see above). Cells were observed under Zeiss Axioplan 2 Imaging Microscope and photos were taken by using Micro-Manager 1.3 software.

### BioLector growth assays

Exponentially growing cells of the strains were collected and 1.5 ml micro-cultures were set at least in triplicate in a 48-well plate starting from an OD_600_ = 0.25. Micro-cultures were left growing in BioLector machine (m2p labs), shaking and under ideal humidity conditions until they reached stationary phase. OD_600_ was monitored every 20 min and recorded electronically. The growth assay was performed twice. Results from the OD measurements of 3 independent micro-cultures of each strain were averaged point-by-point and normalized over the first value of each series before plotting the relative growth rate.

### Drug sensitivity growth assays

Cells were grown in rich medium (YE3S) to an OD_600_∼1. Five serial dilutions (1/10) of cells were spotted on YE3S plates containing drugs at the following concentrations: 10mM hydroxyurea (HU); 30μM camptothecin (CMP); 0.5μM 4-Nitroquinoline N-oxide (4NQO); 0.01% methyl methanesulfonate (MMS); 0.5μg/ml bleocin; or 3.5μg/ml bleomycin. In some growth assays cells were exposed to 100J/m^2^ ultraviolet light (UV-C) immediately after spotting. Plates were incubated at 32°C, except for Bleocin plates which were incubated at 26^°^C, and photos were taken after 2-3 days.

### Flow cytometry

Flow cytometry analyses were performed as previously described (Gregan et al., 2003). Prototrophic strains were arrested in G1 phase by growing for 16h at 26^0^C in EMM medium minus nitrogen, before releasing from the arrest in EMM plus nitrogen.

### Western blot analysis

Proteins were extracted by TCA method and Western blotting was performed as previously described (Ralph et al., 2006). To prepare non-reduced protein samples a modified TCA protein extraction protocol was followed, as previously described (Dardalhon et al., 2012). For the GFP/CFP-tagged proteins the mouse monoclonal antibody Roche anti-GFP (Roche 11814460001) was used. Tubulin was used as a loading control and the probing was done by using mouse monoclonal anti-α-tubulin (Sigma T5168). The secondary anti-mouse antibody used was Peroxidase anti-mouse IgG (Vector Laboratories PI-2000). Detection of the protein signal was done by using Amersham ECL Prime Western Blotting Detection Reagent kit (RPN 2232) and exposing the membrane in an Alpha Innotech FluorChem 8800 Detector.

### Measurement of dNTP levels

dNTP levels were measured as previously described (Moss et al., 2010) with some modifications in the protocol. Briefly, strains were grown to log phase in YE3S and 5 x10^8^ cells were collected by centrifugation and washed once with 3% glucose solution. The cell pellet was kept on ice, while 50μlt of 10% TCA solution were added with brief stirring. Samples were immediately frozen at -80^0^C before proceeding to HPLC analysis. Thawed samples were centrifuged and 20μl of the supernatant was mixed with 80μl of milli Q water (final concentration of 2% TCA). Nucleotides were neutralized by extracting the samples with 100μl of FREON (1,1,2 trichlorotrifluorethane): trioctylamine (4:1). After centrifugation, the supernatant (∼100μl) was collected and 40 μL was injected onto the column for analysis. Chromatography was carried out based on a previously described method on a Waters 2695 system with diode array detection (Waters 2996), monitoring at 254 nm (Moss et al., 2010; Pires et al., 2010). After separation, nucleotides were identified and quantitated against commercially available dNTPs.

### Mutation rate assays

Mutation rates were determined using fluctuation analysis, using mutation to canavanine resistance (Can^R^) (Shcherbakova and Kunkel, 1999) and reversion of an *ade6-M210* mutation (a CCT to CTT mutation causing a P489L mutation) (Catlett and Forsburg, 2003). For the Can^R^ assay, cells were plated onto PMG plates containing 70μg/ml canavanine and were left at 32^°^C for 10 days, before counting resistant colonies, while with Ade^+^ reversion assay, plates were left for 21 days before counting. The mutation rate was calculated with the Ma-Sandri-Sarkar Maximum Likelihood Estimator (MSS-MLE) method, using the Fluctuation AnaLysis CalculatOR (FALCOR) web tool (Hall et al., 2009).

## Acknowledgements

We thank Tony Carr, Oliver Fleck, Christian Holmberg, Tim Humphrey, Olaf Nielsen and Chen-Chun Pai for plasmids and strains. We are grateful to Jürg Bähler for kind permission to use the BioLector technology in his laboratory at UCL and to Lisa Folkes for help with the HPLC analyses at Oxford Institute for Radiation Oncology.

We thank Chen-Chun Pai for comments on the manuscript.

## Author contributions

SEK conceived the research project, CA designed and performed the experiments, CA and SEK wrote the manuscript, IS contributed to the inception of the project and to strain constructions.

---

**Supplementary Figure S1.**
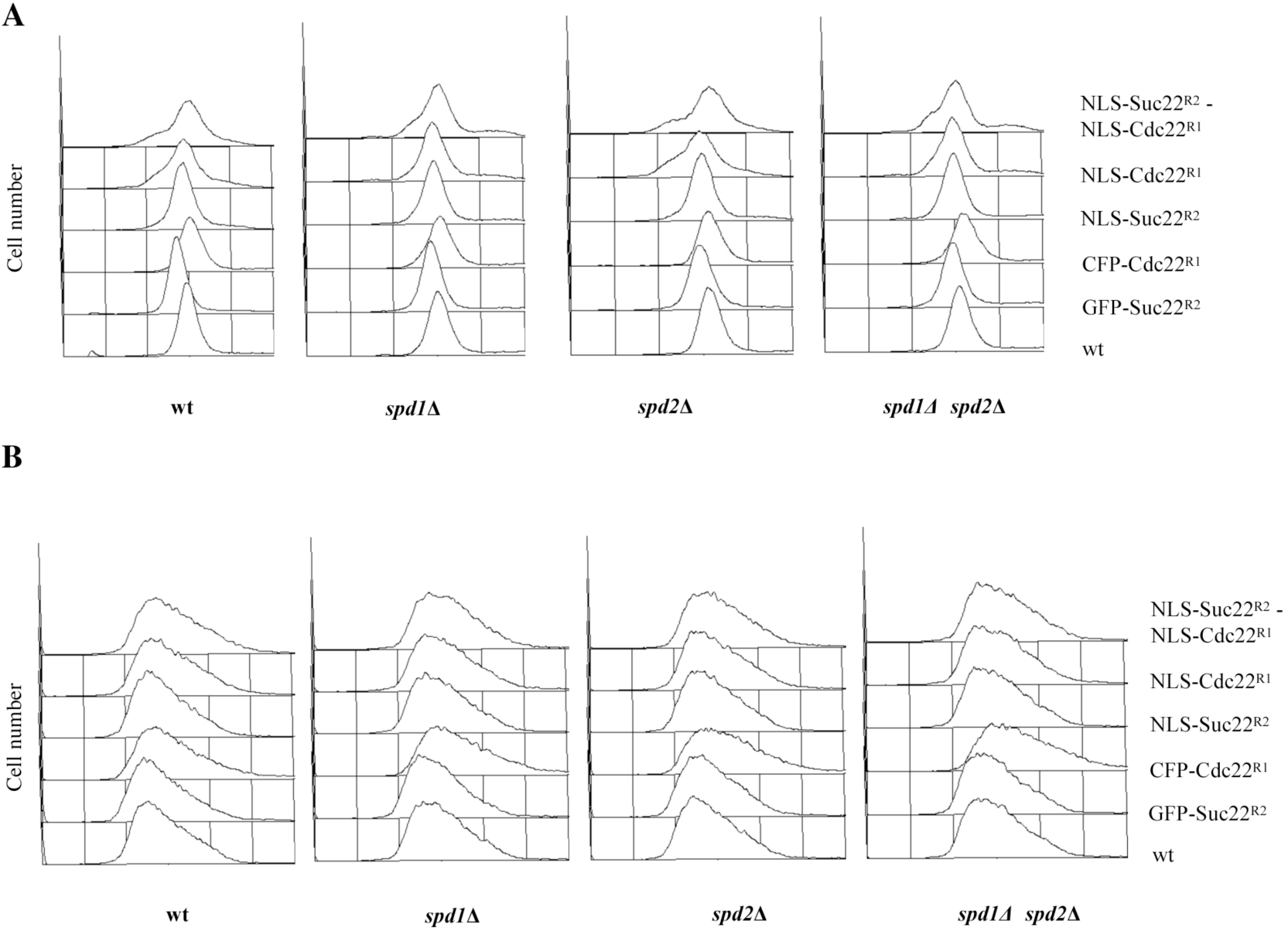
Effect of Spd1 deletion on the S phase defects of RNR mutants. (A) DNA content and (B) Size (forward scatter) of log phase strains after flow cytometric analysis (from bottom to top, strains used were for *spd1*Δ 3367, 3363, 3368, 3365, 3372, 3369; for *spd2*Δ 3399, 3400, 3401, 3402, 3403, 3404; for *spd1*Δ *spd2*Δ 3424, 3429, 3425, 3405, 3414, 3415).

**Supplementary Figure S2.**
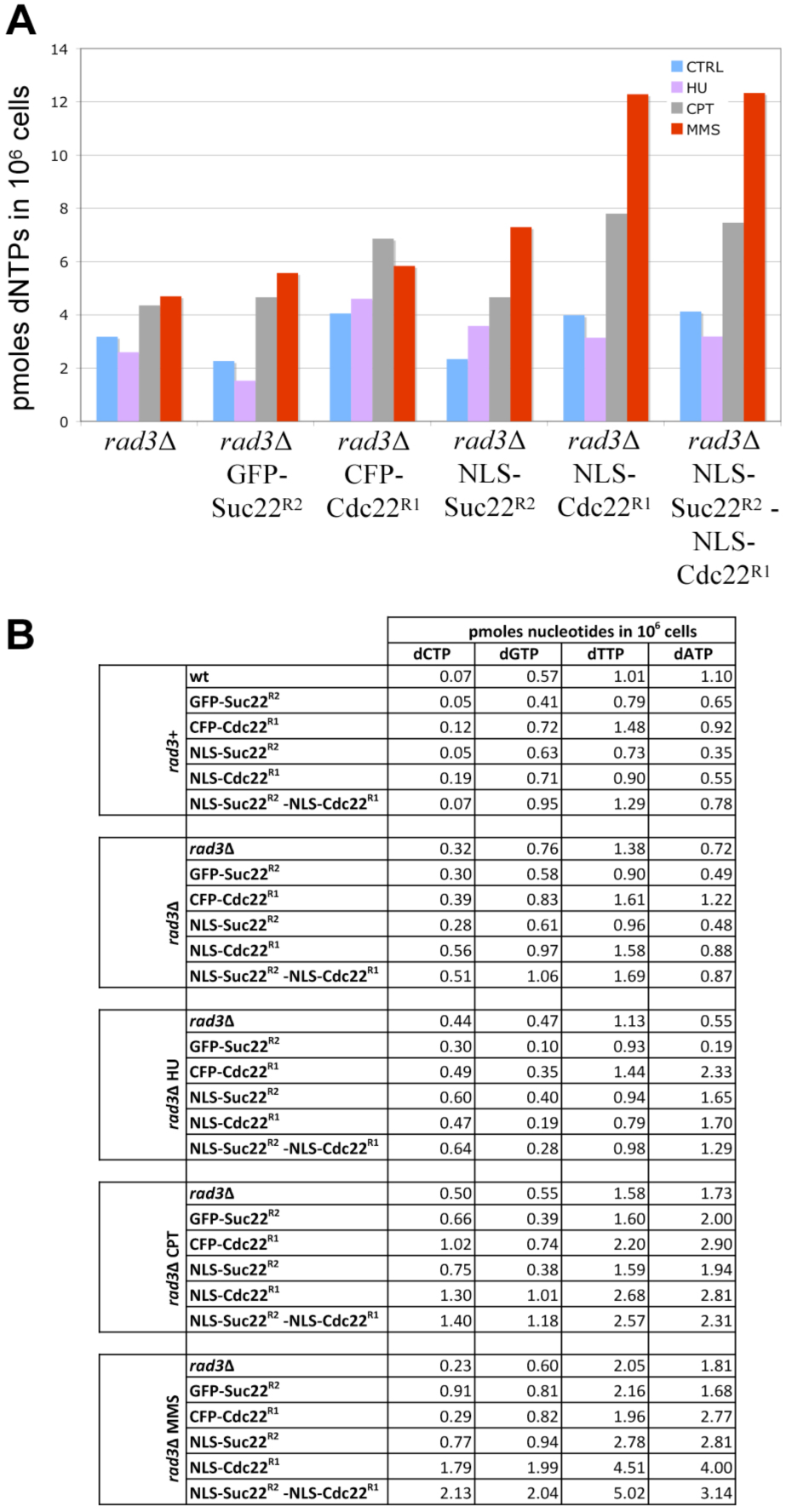
dNTP levels in RNR strains after treatment with DNA damaging agents. (A) Cells were grown in YE3S to exponential phase and treated with 10mM HU for 2hrs, 30μM Camptothecin for 2hrs or 0.01% MMS for 3 hours. dNTP levels were measured by HPLC. Values shown are averages of two or more determinations. *rad3*Δ strains used were 3406, 3423, 3407, 3428, 3408 and 3409 (left to right). (B) The quantity of each nucleotide in the above strains is shown (top to bottom). Note that the amount of nucleotides in a *rad3*+ background is also shown in the top of the table for comparison (strains 138, 2342, 2344, 3167, 3276, 3277; top to bottom).

**Supplementary Table S1.** Strains used in the study.

| Strain | Genotype |
| --- | --- |
| 138 | h- *ade6* M210 *leu1-32 ura4-D18* |
| 2342 | h- GFP-Suc22 *ade6-704 leu1-32 ura4-D18* |
| 2344 | h+ CFP-Cdc22::KanMX6 *ade6-704 leu1-32 ura4-D18* |
| 3167 | h- NLS-GFP-Suc22::natMX6 *ade6-M210 leu1-32 ura4-D18* |
| 3276 | h- NLS-GFP-Cdc22::kanMX6 *ade6-M210 leu1-32 ura4-D18* |
| 3277 | h- NLS-GFP-Suc22::natMX6 NLS-GFP-Cdc22::kanMX6 *ade6-M210 leu1-32 ura4-D18* |
| 2299 | h- 972 |
| 2300 | h+ 975 |
| 3549 | h- GFP-Suc22 |
| 3164 | h+ CFP-Cdc22::KanMX6 |
| 3327 | h- NLS-GFP-Suc22::natMX6 |
| 3328 | h- NLS-GFP-Cdc22::kanMX6 |
| 3329 | h- NLS-GFP-Suc22::natMX6 NLS-GFP-Cdc22::kanMX6 |
| 3406 | h- *rad3Δ*::URA4 h- *ade6* M210 *leu1-32 ura4-D18* |
| 3423 | h- *rad3Δ*::URA4 GFP-Suc22 *ade6-704 leu1-32 ura4-D18* |
| 3407 | h+ *rad3Δ*::URA4 CFP-Cdc22::KanMX6 *ade6-704 leu1-32 ura4-D18* |
| 3428 | h- *rad3Δ*::URA4 NLS-GFP-Suc22::natMX6 *ade6-M210 leu1-32 ura4-D18* |
| 3408 | h- *rad3Δ*::URA4 NLS-GFP-Cdc22::kanMX6 *ade6-M210 leu1-32 ura4-D18* |
| 3409 | h- *rad3Δ*::URA4 NLS-GFP-Suc22::natMX6 NLS-GFP-Cdc22::kanMX6 *ade6-M210 leu1-32 ura4-D18* |
| 3602 | *rad3Δ*::URA4 |
| 3620 | h- *rad3Δ*::URA4 NLS-GFP-Suc22::natMX6 |
| 3584 | h- *rad3Δ*::URA4 NLS-GFP-Cdc22::kanMX6 |
| 3613 | h- *rad3Δ*::URA4 NLS-GFP-Suc22::natMX6 NLS-GFP-Cdc22::kanMX6 |
| 3367 | h- *spd1Δ*::URA4 *ade6* M210 *leu1-32 ura4-D18* |
| 3363 | h- *spd1Δ*::URA4 GFP-Suc22 *ade6-704 leu1-32 ura4-D18* |
| 3368 | h+ *spd1Δ*::URA4 CFP-Cdc22::KanMX6 *ade6-704 leu1-32 ura4-D18* |
| 3365 | h- *spd1Δ*::URA4 NLS-GFP-Suc22::natMX6 *ade6-M210 leu1-32 ura4-D18* |
| 3372 | h- *spd1*Δ::URA4 NLS-GFP-Cdc22::kanMX6 *ade6-M210 leu1-32 ura4-D18* |
| 3369 | h- *spd1*Δ::URA4 NLS-GFP-Suc22::natMX6 NLS-GFP-Cdc22::kanMX6 *ade6-M210 leu1-32 ura4-D18* |
| 3399 | h- *spd2*Δ::hyg+ *ade6 M210 leu1-32 ura4-D18* |
| 3400 | h- *spd2*Δ::hyg+ GFP-Suc22 *ade6-704 leu1-32 ura4-D18* |
| 3401 | h+ *spd2*Δ::hyg+ CFP-Cdc22::KanMX6 *ade6-704 leu1-32 ura4-D18* |
| 3402 | h- *spd2*Δ::hyg+ NLS-GFP-Suc22::natMX6 *ade6-M210 leu1-32 ura4-D18* |
| 3403 | h- *spd2*Δ::hyg+ NLS-GFP-Cdc22::kanMX6 *ade6-M210 leu1-32 ura4-D18* |
| 3404 | h- *spd2*Δ::hyg+ NLS-GFP-Suc22::natMX6 NLS-GFP-Cdc22::kanMX6 *ade6-M210 leu1-32 ura4-D18* |
| 3424 | h- *spd1*Δ::URA4 *spd2*Δ::hyg+ *ade6 M210 leu1-32 ura4-D18* |
| 3429 | h- *spd1*Δ::URA4 *spd2*Δ::hyg+ GFP-Suc22 *ade6-704 leu1-32 ura4-D18* |
| 3425 | h+ *spd1*Δ::URA4 *spd2*Δ::hyg+ CFP-Cdc22::KanMX6 *ade6-704 leu1-32 ura4-D18* |
| 3405 | h- *spd1*Δ::URA4 *spd2*Δ::hyg+ NLS-GFP-Suc22::natMX6 *ade6-M210 leu1-32 ura4-D18* |
| 3414 | h- *spd1*Δ::URA4 *spd2*Δ::hyg+ NLS-GFP-Cdc22::kanMX6 *ade6-M210 leu1-32 ura4-D18* |
| 3415 | h- *spd1*Δ::URA4 *spd2*Δ::hyg+ NLS-GFP-Suc22::natMX6 NLS-GFP-Cdc22::kanMX6 *ade6-M210 leu1-32 ura4-D18* |
| 3285 | h+ Cdc22-D57N |
| 3576 | Cdc22-D57N (prototrophic) |
| 3581 | h- NLS-GFP-Cdc22::kanMX6 D57N |

**Supplementary Table S2.**
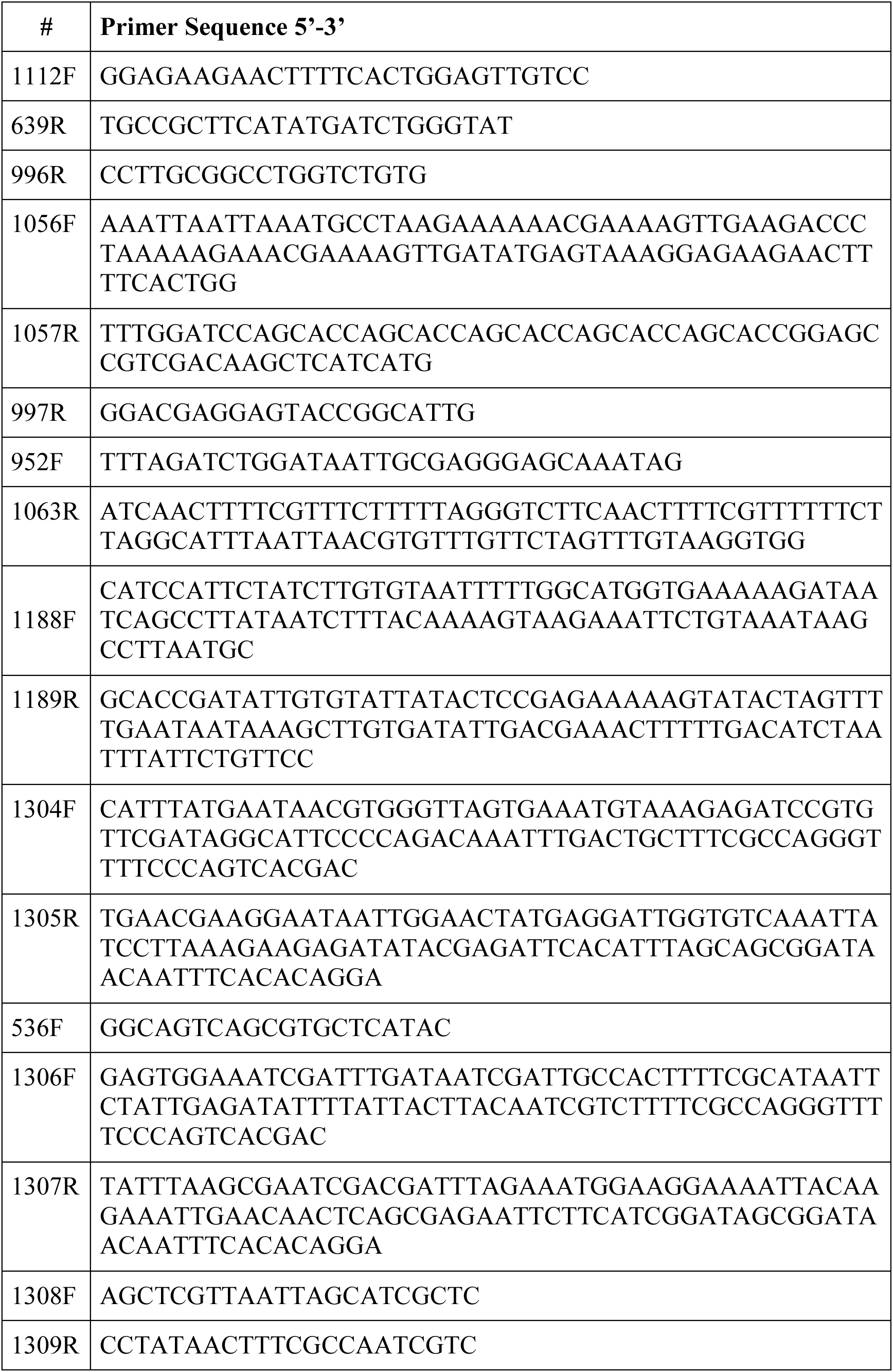
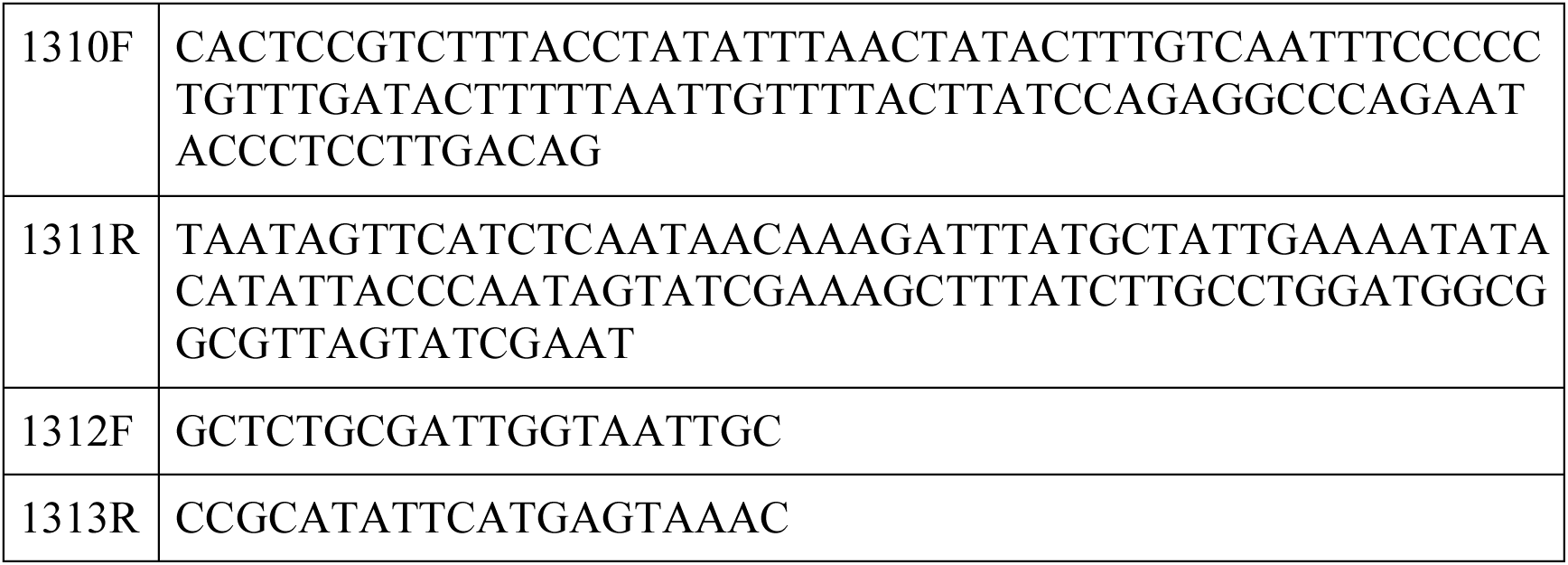
Primers used in the study.

